# On-chip non-contact mechanical cell stimulation - quantification of SKOV-3 alignment to suspended microstructures

**DOI:** 10.1101/2024.02.28.582630

**Authors:** Sevgi Onal, Maan M. Alkaisi, Volker Nock

## Abstract

Although the accumulation of random genetic mutations have been traditionally viewed as the main cause of cancer progression, altered mechanobiological profiles of the cells and microenvironment also play a major role as a mutation-independent element. To probe the latter, we have previously reported a microfluidic cell-culture platform with an integrated flexible actuator and its application for sequential cyclic compression of cancer cells. The platform is composed of a control microchannel in a top layer for introducing external pressure, and a polydimethylsiloxane (PDMS) membrane from which a monolithically-integrated actuator protrudes downwards into a cell-culture microchannel. When actively actuated, the integrated actuator, referred to as micro-piston, transfers the pressure from the control channel as a mechanical force to the cells underneath. When not actuated, the micro-piston remains suspended above cells, separated from the latter via a liquid-filled gap of ∼108 µm. Despite the lack of direct physical contact between the micro-piston and cells in the latter arrangement, we observed distinct alignment of SKOV-3 ovarian cancer cells to the piston shape. To characterize this observation, micro-piston localization, shape, and size were adjusted and the directionality of a mono-layer of SKOV-3 cells relative to the suspended structure was probed. Cell alignment analysis was performed in a novel, label-free approach by measuring elongation angles of whole cell bodies with respect to micro-piston peripheries. Alignment of SKOV-3 cells to the structure outline was significant for circular, triangular and square micro-piston when compared to control areas without micro-piston on the same chip. The effect was present irrespective of whether cells were loaded with micro-pistons in static position (∼108 µm gap) or actively retracted using vacuum (>108 µm gap). Similar alignment was not observed for MCF7 cancer cells and MCF10A non-cancerous epithelial cells. The reported observation of directional movement and growth of SKOV-3 cells towards the region under micro-pistons point towards a to-date unexplored mechanotactic behaviour of these cells, warranting future investigations regarding the mechanisms involved and the role these may play in cancer.

## 1. Introduction

Biophysical cues and mechanical forces are known to direct cell behaviour and tissue development [1, 2, 3, 4]. They play key roles in cellular signalling [5], direct stem cells during development and regeneration [6], and, via tissue stiffness, are related to specific disease characteristics [7]. In vitro, direct mechanical stimulation via geometric cues and solid contact guidance have an impact on cell patterning and directing cell response [8, 9, 10, 11, 12]. However, a potential mechanical cell stimulation by the presence of non-contacting structures in the proximity of cells in a microenvironment has not been investigated so far.

The organization and complexity of the *in vivo* microenvironment are not always fully represented in conventional and bulk cell-culture systems used for *in vitro* studies. Compared to these systems, microfluidics-based biomechanical microdevices can provide extensive spatial and temporal control in both 2D and 3D settings. They can further be designed to introduce static and dynamic physical inputs, and gradients in a physiologically relevant manner [13]. Such microdevices also enable real-time monitoring or visualization. However, cell culture and response in microfluidic devices is not always straightforward unless device design and operation are optimized. For the work presented here, dimensions and features of our so-called micro-piston device were designed based on mammalian cell-culture standards, which encompassed material and structural selections [14]. After characterization of the fabricated devices, the platform was then evaluated for cell-culture, growth, and static mechanical stimulation, by running control and test experiments on a single device.

In addition to its dynamic compression capability, which was reported in our previous work [14, 15], the micro-piston device can be used simply in its static condition by employing the micro-piston as a hanging structure separated from the cells via a liquid-filled vertical gap. Typically, the device remains in this configuration during cell loading and mono-layer formation in preparation for compression studies. During such preparations for our previous work, it was observed that individual SKOV-3 ovarian cancer cells showed distinct alignment to the micro-piston outline. This happened despite the underside of the micro-piston being suspended above the spread cells at distances >30x their height when spread. Due to the absence of reports of such behaviour in literature, this was further investigated using the abilitiy of our platform to provide different micro-piston geometries (i.e. shapes and sizes). By modifying the micro-piston shape, the spatial control on-chip for patterning and redirection of cells was extended while their behavior was influenced by the physical input of the hanging structure in the microchannels. In this study we report the results of these observations, which support the hypothesis of the existence of a previously undocumented cell alignment mechanism in SKOV-3 cells under static conditions. Our results illustrate that suspended structures, such as the micro-pistons found in our platform, can be used to investigate on-chip static mechanical stimulation, thus providing new insights into cell biomechanics in cancer microenvironments.

Parameters such as cell morphology, alignment, elongation, and growth are all affected by mechanical stimulation. While the overall purpose of the original microdevice was to provide dynamic compression, interesting effects of the non-actuated structure on cell-culture were observed after cell loading and growth [16]. In the following we describe how cell alignment on our platform was analyzed in a label-free approach via morphological changes, implemented for this work by measuring elongation angles of cells with respect to micro-piston peripheries. This novel approach relies on the orientation of whole cell bodies rather than subcellular parts such as nuclei alone. Using this, we compare the directional movement and growth of SKOV-3 cells in the region under micro-pistons to control areas on respective devices. While we observed that individual behaviour of SKOV3 cells is comparable to the collective behaviour of MCF7 cancer cells and MCF10A non-cancerous epithelial cells in micro-piston devices, the latter two cell types did not show quantifiable alignment. Cell growth and spread under the micro-pistons were investigated for different piston shapes including circular, square, and triangular, and piston sizes, as well as for different cell loading conditions, proposing that SKOV-3 cell movements were influenced by the presence of the suspended, static micro-pistons.

## 2. Materials and methods

For a comprehensive description of the design, fabrication, and use of the cell-compression microdevice the reader is referred to our previous work [14]. The following provides a summary of the key differences related to fabrication, operation and characterization between this and the current work.

### 2.1. Device design and fabrication

Briefly, the device consists of a top layer forming the control channel, which is bonded to a middle layer with flexible membrane and monolithically attached micro-piston. This in turn is suspended into a bottom microchannel, all made of polydimethylsiloxane (PDMS) elastomer. Finally, this PDMS construct is enclosed by a glass substrate to form a chamber for cell-culture and mechanical stimulation (Fig. 1(a)). The fabrication of the PDMS layers included different soft-lithography methods, with the top control layer fabricated using standard replica molding, while the relatively thinner membrane/micro-piston layer was achieved by spin-coating of the PDMS on a Si master. The bottom layer containing top-and-bottom open microchannels was realized via exclusion molding of PDMS from a separate Si master. The PDMS formulation used for all layer fabrications was 10:1 w/w (Sylgard 184, Dow) base and curing agent. Curing conditions for each layer, spin-coating settings for PDMS membrane/micro-piston layer and the exclusion molding setup for the bottom layer were as detailed in [14]. After the individual PDMS layers were ready, these were aligned and plasma-bonded to each other by manually matching the alignment marks in each layer. Finally, the completed PDMS assembly was then plasma-bonded to a glass slide. Different diameters and shapes including the circular, triangle and square micro-pistons shown in Fig. 1(a) were fabricated in the same manner.

**Figure 1:**
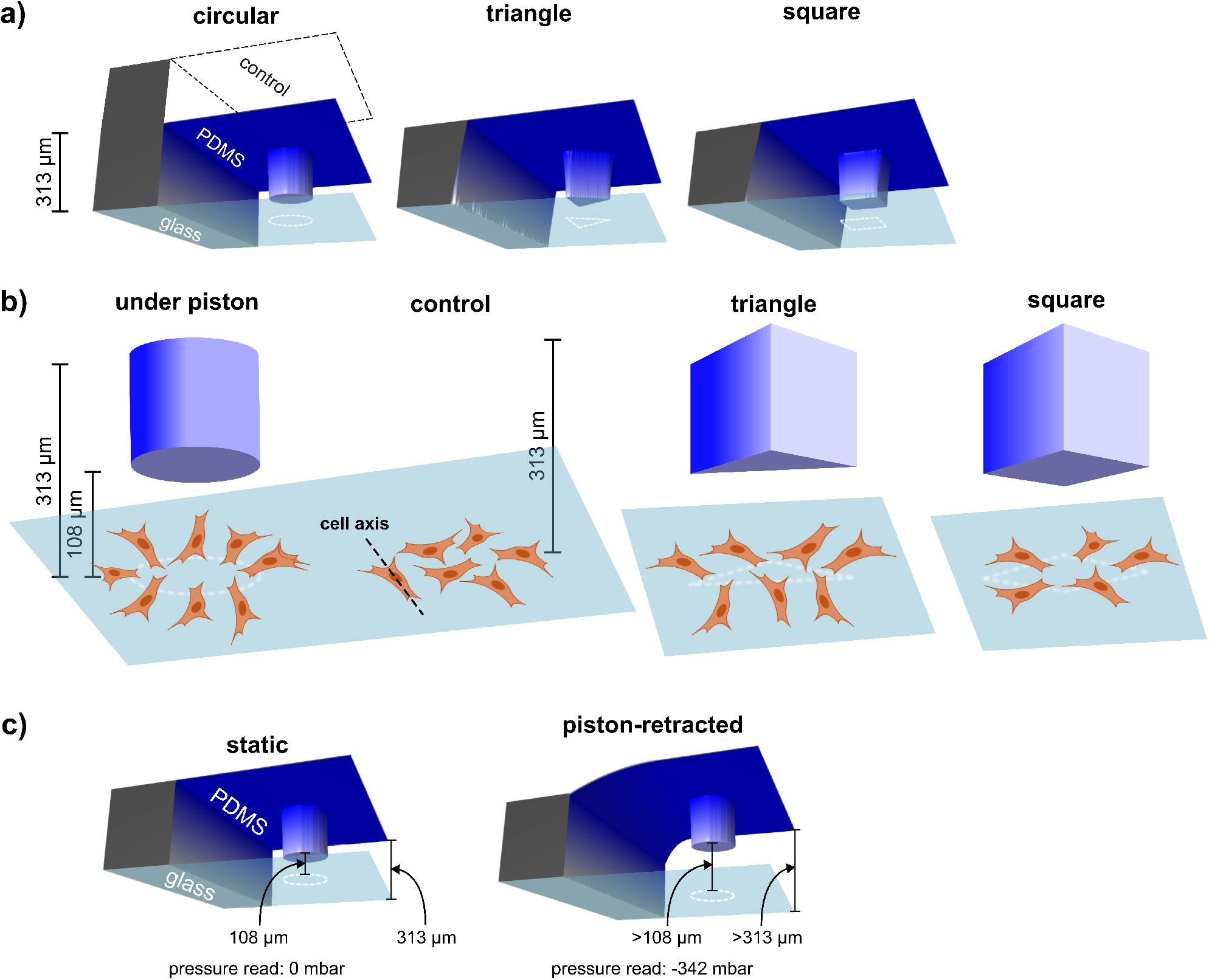
Schematic showing devices fabricated with different micro-pistons, principle of cell alignment under micro-pistons and in control regions, and observed cell response as a function of different micro-piston shapes. (a) Composite images of actual 3D optical profilometer scans of the fabricated PDMS devices showing circular, triangle and square micro-pistons attached to membranes and suspended into microchannels formed by the bottom layers. Control channel and glass layers are also indicated to illustrate device operation. (b) Illustration of the method used to test cell alignment in this study. Alignment with respect to the periphery of a micro-piston suspended at a gap height of on average 108 µm was compared to random cell movement taking place in a control region in the same microchannel (height of on average 313 µm) via the main cell axis. This was done for circular, triangle and square micro-pistons acting as suspended structures. (c) Illustration of the two cell loading methods compared in this work. In the static case, no actuation was applied (0 mbar), while for the piston-retracted case, the membrane and piston were lifted to >108 µm above the cell layer via application of vacuum (−342 mbar) to the control channel.

### 2.2. Optical profilometer measurements

All fabricated micro-piston devices were scanned with 3D optical profilometry (Profilm3D, Filmetrics) to ensure experimental repeatability and characterize the distance between the bottom of the micro-piston and the cells (Fig. 1(a)). Data for the dimensions of the device compartments were processed and analyzed using the ProfilmOnline software. Using this approach, as was also characterized in [14], the height of the cell culture channel and micro-piston of PDMS replicas were measured to be on average 313 µm and 205 µm, respectively (Fig. 1(b)). After the device layers were assembled, the resultant gap between the bottom of the micro-piston and the glass surface was measured to be on average 108 µm (Fig. 1(b)).

### 2.3. Cell culture and preparation

SKOV-3 ovarian cancer cells (kindly supplied by the Laboratory for Cell and Protein Regulation at the University of Otago, New Zealand) were cultured in Earle’s salts and L Glutamine positive MEM medium (GIBCO^*®*^) supplemented with 10% fetal bovine serum (FBS, Life Technologies), 1% of penicillin/ streptomycin (Life Technologies) and 0.2% fungizone (Life Technologies) in a humidified atmosphere of 5% CO_2_ at 37°C. Detailed cell-culture media components and concentrations are provided in Supplementary Table S1 for SKOV-3 and other two cell types used, MCF7 and MCF10A (kindly supplied by Dr Tracy Hale from Massey University, New Zealand). Initial cell seeding for the three cell types was tested with concentrations of 0.5, 1, 1.5 and 1.8M cells/ml to optimize the cell number in microfluidic channel for formation of a cell mono-layer and to capture sufficient cells under the micro-piston. Based on these tests, all experiments were performed with 1.5M cells/ml as the optimized cell concentration.

### 2.4. Device operation for cell culture and mechanical stimulation experiments

Before any cell-culture on-chip was commenced, micro-piston devices were sterilized under UV light in a biological safety cabinet for at least 30 minutes. Cells were introduced into the culture channels while the micro-piston was either in static state (no pressure applied to the control channel) or further driven-up (retracted) towards the control channel (Fig. 1(c)). For the piston-retracted loading method, the micro-pistons were retracted by applying negative pressure to the control channel. This was achieved by removing 2 cm^3^ of air over 4 minutes using a syringe pumping system, which corresponded to -342 mbar of pressure read in the control channel. When loading this way, the micro-piston was kept retracted for 5 minutes after cell loading to let the cells sink to the poly-L-lysine (PLL)-coated glass surface [14] of the bottom microchannel. Then the micro-piston was released to recover its initial position by applying positive (increasing) pressure via the control. For either case, cells were cultured in the micro-piston devices for at least 2 days, and their growth and spread around and under the micro-pistons were then monitored and recorded during culture to observe the impact of of the micro-piston on cell arrangement.

For all samples, cells were kept in culture media to eliminate any non-mechanical stress and the microscopy focal plane was kept on the cells while imaging. Progression of cell culture and mechanical response of the cells were recorded with time-lapse imaging.

Since microchannels in flexible microdevices contain very low liquid volumes compared to conventional cell-culture containers, such as flasks or plates, evaporation of channel contents can be a problem when devices are placed in an incubator at 37°C. In the current work, excess evaporation was prevented by closure of inlets and outlets, use of a humidity chamber and by gently replenishing media every 1 or 2 days during cell-culture with the same type of growth media.

### 2.5. Imaging and data analysis

Images of cells were taken as phase-contrast image series using an inverted microscope (Nikon ECLIPSE Ts2) equipped with a digital camera (Tucsen USB2.0 H Series) and TCapture imaging software. For cell growth, alignment and spread analysis, cells were imaged day-to-day starting from the cell seeding time using 10X (NA 0.25; Ph1 ADL) and 20X (NA 0.45; Plan-Fluor OFN22 Ph1 ADM) objectives.

Furthermore, bright-field microscopy live-cell image sequences of SKOV-3 cancer cells proliferating inside the cell chamber and orienting under the periphery of the micro-piston were recorded over a period of 48 hours. Images were collected every 30 minutes in an incubator of 37°C and 5% CO_2_ using a live cell imaging system (CytoSmart). Automated analysis of area coverage (CytoSmart) was also applied.

Recorded images were processed and analyzed using ImageJ (Fiji) [17]. Cell orientation under the micro-pistons and in control areas (Fig. 1(b)) was analyzed from phase contrast images in a label-free approach. During image analysis, ROIs were fitted around each micro-piston and cells that were in part under a micro-piston were counted towards the analysis. For cell alignment analysis, the ROIs of the cells that elongated across the “virtual” (as projected) periphery of a micro-piston (Fig. 1(b)) were outlined via the freehand tool of ImageJ. The cell elongation angle calculation method is illustrated in Figure 2 for the different micro-piston shapes. The x-y positions of cells were measured from the ROIs of the cell outlines and recalculated relative to the centroid of the micro-pistons. Angles of cells were measured using ImageJ as the angle between the major axis of the ellipse fitted on the cell ROI and a line parallel to the field of view (FoV). The center of this ellipse is the x-y coordinates of the centroid of each cell. Thus, the angle of the cell line passing through the centroid of the cell ROI and the centroid of the piston could be measured via the arctan of the x-y position of a cell relative to the x-axis of the FoV. For the cell elongation angle with respect to the circular micro-piston periphery, the angle of the cell (calculated relative to the x-axis of the FoV) was recalculated relative to the tangent at the circular micro-piston perimeter by taking into account the difference with the angle of the cell line.

**Figure 2:**
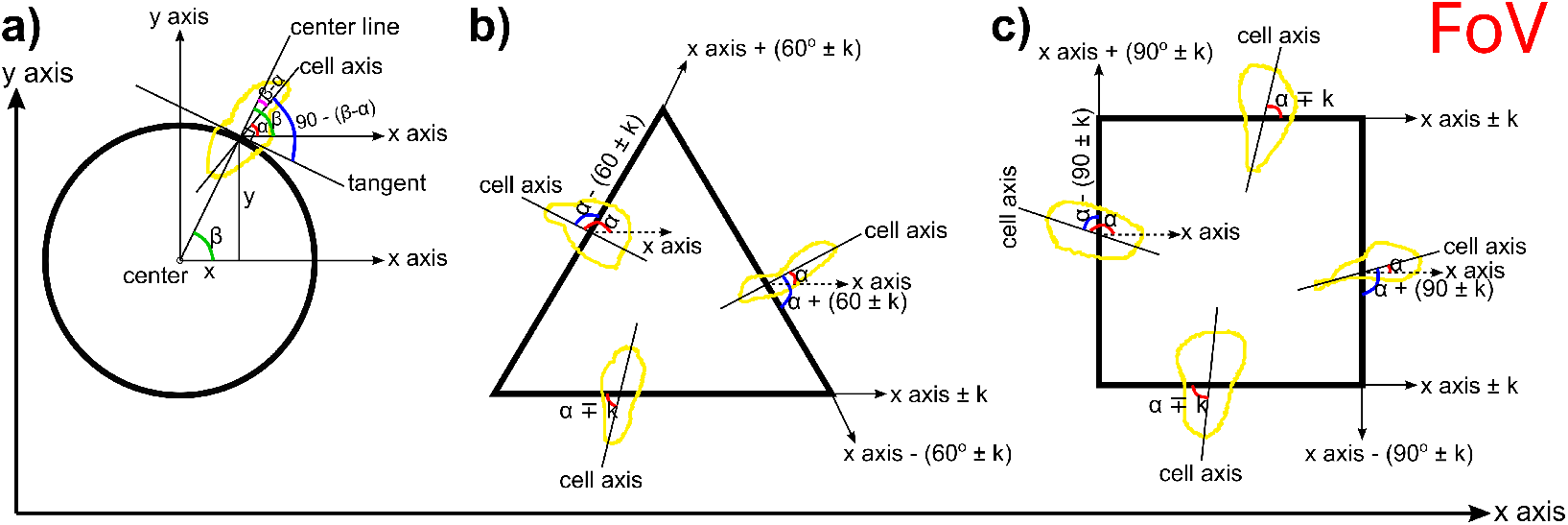
Cell elongation angle measurement and calculation method. Cells elongating across the periphery of micro-pistons were measured and quantified for their elongation angle relative to the perimeter of each micro-piston. (a) Cell elongation angle at the periphery of a circular micro-piston was calculated as given in Eqn. 1. The centroids of cell shapes were calculated as (x,y) of the cell centroid relative to the (x,y) of the piston centroid and this information gave the cell position measured as (x_cell_, y_cell_). These cells were the ones elongating across the periphery of circular pistons. Their coordinates were used in the calculation of the cell elongation angle with respect to piston perimeters. Cell axis was taken as the major axis of the ellipse fitting the cell shape, passing from the cell centroid. The cell angle ‘α’ was determined by the angle of the cell axis relative to the x-axis of the FoV (see Eqn. 2). Then cell angle relative to the tangent was calculated based on the arctan angle ‘β’ of the center line passing from the cell centroid coordinates given here as cell position (xcell, ycell) (see Eqn. 3) and cell angle ‘α’. (b) Cell elongation angles at the periphery of a triangular micro-piston were measured relative to the x-axis of the FoV as ‘α’ and re-calculated relative to the corresponding edge angles, which were also measured relative to the x-axis of the FoV and formulated as based on the expected angle of 60 degrees and correction angle ‘k’. (c) Cell elongation angles at the periphery of a square micro-piston were measured relative to the x-axis of the FoV as ‘α’ and re-calculated relative to the edge angles, which were also measured relative to the x-axis of the FoV and formulated as based on the expected angle of 90 degrees and correction angle ‘k’.

Cell elongation angle at the periphery of a circular micro-piston was calculated as given below:

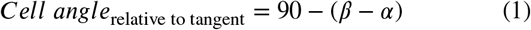

where,

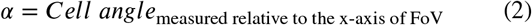

and, for the cell position measured as (x_cell_, y_cell_),

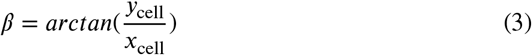

Cell elongation angles at the periphery of triangular and square micro-pistons were measured relative to the x-axis of the FoV and re-calculated relative to the angles of the corresponding edges for each shape, which were also measured relative to the x-axis of the FoV. Representative geometries of micro-pistons overlaid onto random areas of the cell mono-layer around the micro-piston were used as control. Cell alignment and elongation angle analysis in those geometries was performed in the same manner as under micro-piston peripheries. Thus, cells under the periphery of micro-pistons formed the test group, while cells around micro-pistons provided a control group in the same micro-channel. To Two sample t-test between percents (StatPac) was used to compare percentages of individual groups in cell elongation angle data. χ^2^ test was used to compare distributions of cell elongation angle data by the frequencies of angle types in each group. Statistical significance was taken as p < 0.05.

## 3. Results and discussion

It is well known that cells exist in a complex biochemical and mechanical equilibrium with their surrounding microenvironment and arrange themselves relative to this [18]. When cultured *in vitro*, the mechanical component of this microenvironment includes a variety of engineered structures such as for example the natural or engineered topography of the artificial substrate used. Culture surfaces are regularly fabricated to include micro- and nanostructures [19, 20] with the aim to study and influence cell behaviour via direct mechanical interaction [21, 22, 23]. With more and more cell-culture studies now performed in microfluidic devices [24], cells encounter additional structures, besides the mainly culture-surface based ones, for example in the form of channel walls and lids. While physically removed from the cells, these structures are typically still within their relative vicinity due to the micro-scale dimensions of the device features. Besides purely static structures, microfluidic devices can also contain moving ones in the form of micro-valves, variations thereof which can be deliberately suspended close to cells and brought into contact with them to enable active mechanical cell stimulation [14, 15, 16]. Whereas such direct interactions have been studied extensively [13], little is known about how cells respond to suspended structures before physical contact is made. This is in particular the case for when these structures are still at rest and thus are not imparting forces on the cells indirectly via displacement of the liquid in between.

The current work was motivated by observations of SKOV-3 cancer cells exhibiting distinct directional movement and growth underneath micro-pistons when cultured in static media on our sequential compression platform [14, 16]. This indicated a mechanical stimulation by the suspended micro-pistons even though these were not actuated or in physical contact with the cells. To investigate this further, cancer cell growth and elongation in the microfluidic platform were characterised by running control and test experiments on a single device. As illustrated in Fig. 3(a), once loading was complete, directional movement and growth of SKOV-3 cancer cells showed distinct alignment to the shape of the suspended micro-piston outline. The cell number under micro-piston and in the channel increased on average by 80% on Day 2, providing a confluent cancer cell mono-layer on-chip (see Supplementary Movie S1). Thus, cells grew under and around the static micro-pistons regardless of the height difference between the gap under micro-piston and the rest of the channel.

As shown in Figure 3(a), to further study this, cancer cell growth and spread under the micro-pistons were compared to that in corresponding control regions. This was also done for different shapes, such as circular, square, and triangular as well as for varying diameters of the circular shape (Fig. 4(a)), to probe for any shape influence. Starting from Day 1 of culture, all micro-piston shapes exhibited distinct alignment of cell axes with respect to micro-piston perimeter (Fig. 3(a-b) and Fig. 4(a-b)). This was the case despite the significant gap between the SKOV-3 cells, typically extending to a height of around 3 µm when non-confluent [25], and the bottom of the micro-piston suspended on average 108 µm above the cells in static state. The regular alignment of cells under the periphery of the micro-pistons, in the absence of direct contact, showed a clear pattern, matching the size and shape of the projected outline of the bottom of the micro-pistons suspended into the culture channels from above.

**Figure 3:**
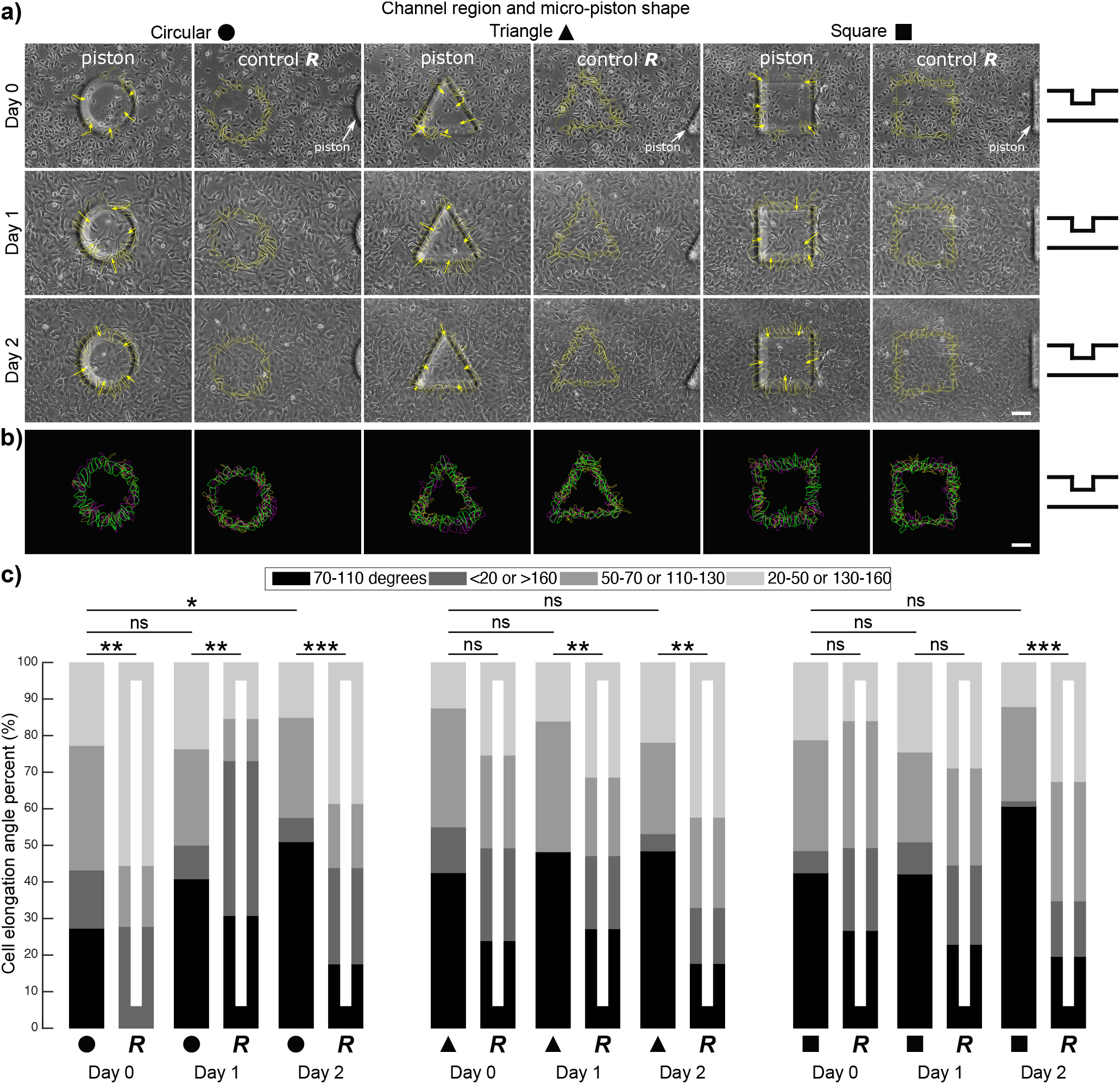
Cancer cell growth, alignment and elongation with respect to micro-piston perimeters of different shapes and in random areas of the corresponding geometries as control. (a) Representative images for SKOV-3 cancer cells occupying the area under 300-µm-diameter circular, triangular and square micro-pistons and in control regions of the microchannel. Cells aligned across the periphery of and grew under the micro-pistons (Scale bar 100 µm). Arrows illustrate the main direction of cell elongation in relation to the suspended micro-pistons. The micro-piston schematic next to the images indicates that micro-pistons were in static state over days (Day 0, 1, and 2). As control for cell alignment, ROIs corresponding to the same micro-piston geometry were used to outline cells in random areas of the cell culture chamber. (b) Cell alignment profiles under the micro-piston periphery and in the control region, showing overlaid outlines of cells over days with respect to the micro-piston shape being circular, triangular or square, color-coded as yellow for Day 0, magenta for Day 1, green for Day 2 (Scale bar 100 µm). Outlines of cells over days are also shown separately for each colour in Supplementary Fig. S1. (c) Cell elongation angles calculated relative to the micro-piston periphery or corresponding geometry as control. On a 0-180 degree scale, angles between 70 and 110 degrees were counted as along the radial elongation across the micro-piston periphery (or corresponding control geometry) and angles of more than 160 and less than 20 degrees were counted as along the micro-piston periphery (or corresponding control geometry). Other angles were grouped as between 50-70 or 110-130 degrees, and between 20-50 or 130-160 degrees. The frequencies of angle groups are shown in percentages over days (Day 0, 1, and 2) for circular, triangular, and square micro-pistons and the corresponding random/control area (*R*) of each geometry. On the plot, n = 44, 36, 76, 52, 106, 57, 40, 67, 56, 70, 64, 85, 33, 75, 57, 83, 66, 92 cells, respectively. Indicators on the horizontal bars show significances of differences between cell elongation angle distribution in the groups (χ^2^ test), *: p < 0.05, **: p < 0.0005, ***: p < 0.0001, ns: not significant.

**Figure 4:**
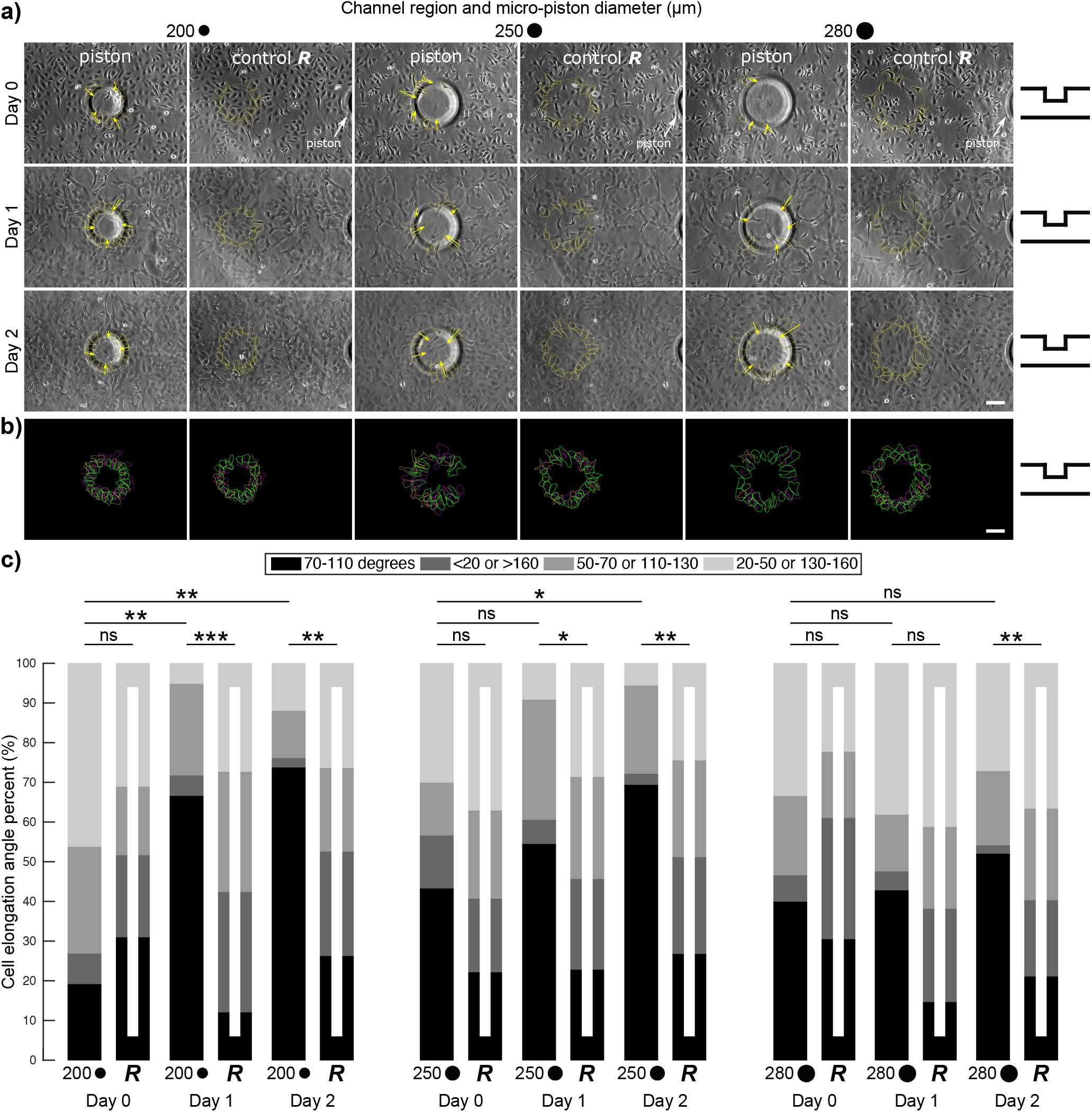
Cancer cell growth, alignment and elongation with respect to micro-piston perimeters of different diameters and in random areas of the corresponding geometries as control. (a) Representative images for SKOV-3 cancer cells occupying the area under circular micro-pistons of different diameters (Scale bar 100 µm). Cells aligned across the periphery of and grew under micro-pistons. Arrows illustrate the main direction of cell elongation in relation to the suspended micro-pistons. The micro-piston schematic next to the images indicates that micro-pistons were in static state over days (Day 0, 1, and 2). (b) Cell alignment profiles under the micro-piston periphery showing merged cell outlines from over days with respect to the circular micro-piston at 200, 250, and 280 µm diameter, color-coded as yellow for Day 0, magenta for Day 1, green for Day 2. (c) Cell elongation angles calculated relative to the circular micro-piston periphery or corresponding diameter as control. On a 0-180 degree scale, angles between 70 and 110 degrees were counted as along the radial direction across the micro-piston periphery (or corresponding control geometry) and angles of more than 160 and less than 20 were counted as along the micro-piston periphery (or corresponding control geometry). Other angles were grouped as between 50-70 or 110-130 degrees, and between 20-50 or 130-160 degrees. The frequencies of angle groups are shown in percentages over days (Day 0, 1, and 2) for different diameters of circular micro-piston of 200, 250, and 280 µm and their corresponding random/control area (*R*) of 200R, 250R and 280R, respectively. On the plot, n = 26, 29, 39, 33, 42, 38, 30, 27, 33, 35, 36, 41, 15, 36, 21, 34, 48, 52 cells, respectively. Indicators on horizontal bars show significances of differences between cell elongation angle distribution in the groups (χ^2^ test), *: p < 0.05, **: p < 0.003, ***: p < 0.0001, ns: not significant.

To the best of our knowledge, similar directional behavior of SKOV-3 ovarian cancer or other cells to hanging physical structures in microchannels, where no direct contact other than through liquid media exists, has not previously been observed. Hsieh *et al*. also reported cell alignment in a microfluidic setting, as a result of compressive strains, however cells in their case were in direct contact with the compression source via solid hydrogel [9]. In an effort to further characterize this behavior, image analysis was used to quantify cell elongation angles relative to each micro-piston shape. Cells elongating across the micro-piston perimeter, and corresponding geometry acting as control in a randomly chosen area of the cell mono-layer away from the piston, was outlined and measured for their angles. This was achieved mainly by measuring cell angle relative to the x-axis of the FoV and then re-calculating the angle by taking the corresponding micro-piston shape as reference, as illustrated in Fig. 2. On a 0-180 degree scale, angles between 70 and 110 degrees were counted as along the radial direction across the micro-piston periphery (or the corresponding control geometry) and angles of more than 160 and less than 20 were counted as along the micro-piston periphery (or the corresponding control geometry) [9]. Other angles were grouped as between 50-70 or 110-130 degrees, and between 20-50 or 130-160 degrees.

The orientation of the cells on the alignment profiles showed that cells tended to elongate along the radial direction with respect to micro-piston peripheries compared to random control areas (Fig. 3). Differences in cell elongation angle distribution between the groups under the micro-piston periphery and the corresponding random area were statistically significant (Fig. 3(c)). For circular shaped pistons, cell elongation angles under the periphery of the micro-piston were highly significantly different to the random area on all days. For triangular shaped pistons, the difference in cell angle frequencies between the micro-piston periphery and the random area was not statistically significant on Day 0 (p = 0.0627), but highly significant on Days 1 and 2 (p < 0.0005). For square shaped pistons, the difference was not significant on Day 0 (p = 0.1174) and was (barely) not statistically significant on Day 1 (p = 0.0508), but became highly significant on Day 2 (p < 0.0001). The individual angle groups were further compared for micro-pistons and their corresponding random area peripheries, as shown in Table 1.

**Table 1.** Statistical significance values by two sample t-test between individual angle type percentages for each group of under different shape micro-piston and control (random) areas of the cell mono-layer. The frequencies of the angle groups are shown in percentages over days (Day 0, 1, and 2) for circular micro-piston (C-), circular random/control area (CR-), triangle micro-piston (T-), triangle random/control area (TR-), square micro-piston (S-), and square random/control area (SR-).

| Day | Area | Angles (degrees) |  |  |  | Cells | p-value |  |  |  |
| --- | --- | --- | --- | --- | --- | --- | --- | --- | --- | --- |
|  |  | 70-110 (%) | <20 or >160 (%) | 50-70 or 110-130 (%) | 20-50 or 130-160 (%) | n | 70-110 (%) | <20 or >160 (%) | 50-70 or 110-130 (%) | 20-50 or 130-160 (%) |
| 0 | C | 27.27 | 15.91 | 34.09 | 22.73 | 44 | 0.0011 | 0.2005 | 0.082 | 0.0034 |
|  | CR | 0 | 27.78 | 16.67 | 55.56 | 36 |  |  |  |  |
| 1 | C | 40.79 | 9.21 | 26.32 | 23.68 | 76 | 0.2503 | <0.0001 | 0.0433 | 0.2538 |
|  | CR | 30.77 | 42.31 | 11.54 | 15.38 | 52 |  |  |  |  |
| 2 | C | 50.94 | 6.6 | 27.36 | 15.09 | 106 | 0.0001 | 0.0006 | 0.163 | 0.0009 |
|  | CR | 17.54 | 26.32 | 17.54 | 38.6 | 57 |  |  |  |  |
| 0 | T | 42.5 | 12.5 | 32.5 | 12.5 | 40 | 0.0462 | 0.114 | 0.4287 | 0.114 |
|  | TR | 23.88 | 25.37 | 25.37 | 25.37 | 67 |  |  |  |  |
| 1 | T | 48.21 | 0 | 35.71 | 16.07 | 56 | 0.0161 | 0.0005 | 0.0778 | 0.0489 |
|  | TR | 27.14 | 20 | 21.43 | 31.43 | 70 |  |  |  |  |
| 2 | T | 48.44 | 4.69 | 25 | 21.88 | 64 | 0.0001 | 0.0403 | 0.9677 | 0.0097 |
|  | TR | 17.65 | 15.29 | 24.71 | 42.35 | 85 |  |  |  |  |
| 0 | S | 42.42 | 6.06 | 30.3 | 21.21 | 33 | 0.1075 | 0.0392 | 0.6581 | 0.5139 |
|  | SR | 26.67 | 22.67 | 34.67 | 16 | 75 |  |  |  |  |
| 1 | S | 42.11 | 8.77 | 24.56 | 24.56 | 57 | 0.0167 | 0.0446 | 0.7957 | 0.5696 |
|  | SR | 22.89 | 21.69 | 26.51 | 28.92 | 83 |  |  |  |  |
| 2 | S | 60.61 | 1.52 | 25.76 | 12.12 | 66 | <0.0001 | 0.0043 | 0.3544 | 0.0034 |
|  | SR | 19.57 | 15.22 | 32.61 | 32.61 | 92 |  |  |  |  |

From this it was determined that cell angle percentages in the groups of ‘70-110’ degrees, and ‘<20 or >160’ degrees under the piston peripheries were larger and lower, respectively, compared to those in the randomly selected control areas. While the group of ‘50-70 or 110-130’ degrees largely did not show significant changes between the micropiston and random areas, the group of ‘20-50 or 130-160’ degrees had statistically higher percentages in random areas mainly on Day 2 of cultures. Thus, the main difference between the micro-pistons and control areas was that the cells under the micro-piston peripheries preferred to elongate more in radial direction (70-110 degrees) as opposed to along the peripheries (<20 or >160 degrees) or other angle groups. This was not the case in the random areas for corresponding geometries. Further, there was no statistically significant differences in cell angle frequency distribution over the days, except between Days 0 and 2 under the circular micro-piston periphery (p = 0.039) (Fig. 3(c)). This means that cells exhibited elongation along the radial direction (70-110 degrees) angles under the micro-piston peripheries from early stages of the cell-culture on-chip. This result was in agreement with higher percentages of cell elongation angles along the radial direction under the micro-piston peripheries at almost all days and for all different micro-piston shapes (Fig. 3(c) and Table 1).

Interestingly, this was also visible during live-cell imaging of cells, the latter which were observed orienting under the periphery of a square (Supplementary Movie S1 Clip 1) and triangular micro-piston (Supplementary Movie S1 Clip 2) over 2 days of culture with images taken every 30 minutes. When cultured in microchannels without micro-pistons of 313 µm (Supplementary Movie S1 Clip 3) and 200 µm height (Supplementary Movie S1 Clip 4), cells of the same type showed no discernible preference in terms of orientation. In addition, there was also no statistically significant difference in the frequencies of the cell elongation angles under the micro-piston peripheries between different shapes on all days. Given that, in this case, the different geometry micro-pistons acted as control for each other, this consistency showed that cells followed the guidance provided by the particular shape of the micro-pistons.

Apart from the different shapes, the cell elongation angle was also quantified for different diameters of the circular micro-pistons. This was done to investigate whether size of the shape had any effect on the alignment patterns of cells elongating across the micro-piston peripheries. Compared to control regions, cells again showed distinct alignment and elongated significantly along the radial direction with respect to the circular micro-piston peripheries over multiple days at diameters of 200, 250, and 280 µm (Fig. 4 and Table 2), as well as 300 µm (Fig. 3). Thus, SKOV-3 cells also followed the smaller or larger diameter micro-pistons when presented as hanging structures. Again, this was not observed for corresponding control diameters in randomly-chosen control areas of the cell mono-layer where no micro-piston was present.

**Table 2.** Statistical significance values by two sample t-test between individual angle type percentages for each group of under different diameter circular micro-piston and control (random) areas of the cell mono-layer. The frequencies of the angle groups are shown in percentages over days (Day 0, 1, and 2) for different diameters of circular micro-piston of 200, 250, and 280 µm and their corresponding random/control area of 200R, 250R and 280R, respectively.

| Day | Area | Angles (degrees) |  |  |  | Cells | p-value |  |  |  |
| --- | --- | --- | --- | --- | --- | --- | --- | --- | --- | --- |
|  |  | 70-110 (%) | <20 or >160 (%) | 50-70 or 110-130 (%) | 20-50 or 130-160 (%) | n | 70-110 (%) | <20 or >160 (%) | 50-70 or 110-130 (%) | 20-50 or 130-160 (%) |
| 0 | 200 | 19.23 | 7.69 | 26.92 | 46.15 | 26 | 0.3204 | 0.1779 | 0.3894 | 0.2544 |
|  | 200R | 31.03 | 20.69 | 17.24 | 31.03 | 29 |  |  |  |  |
| 1 | 200 | 66.67 | 5.13 | 23.08 | 5.13 | 39 | <0.0001 | 0.0057 | 0.4909 | 0.0113 |
|  | 200R | 12.12 | 30.3 | 30.3 | 27.27 | 33 |  |  |  |  |
| 2 | 200 | 73.81 | 2.38 | 11.9 | 11.9 | 42 | 0.0001 | 0.0027 | 0.2713 | 0.1029 |
|  | 200R | 26.32 | 26.32 | 21.05 | 26.32 | 38 |  |  |  |  |
| 0 | 250 | 43.33 | 13.33 | 13.33 | 30 | 30 | 0.097 | 0.5937 | 0.3821 | 0.5757 |
|  | 250R | 22.22 | 18.52 | 22.22 | 37.04 | 27 |  |  |  |  |
| 1 | 250 | 54.55 | 6.06 | 30.3 | 9.09 | 33 | 0.0091 | 0.0548 | 0.6747 | 0.0452 |
|  | 250R | 22.86 | 22.86 | 25.71 | 28.57 | 35 |  |  |  |  |
| 2 | 250 | 69.44 | 2.78 | 22.22 | 5.56 | 36 | 0.0004 | 0.0085 | 0.823 | 0.0259 |
|  | 250R | 26.83 | 24.39 | 24.39 | 24.39 | 41 |  |  |  |  |
| 0 | 280 | 40 | 6.67 | 20 | 33.33 | 15 | 0.5177 | 0.073 | 0.7774 | 0.4108 |
|  | 280R | 30.56 | 30.56 | 16.67 | 22.22 | 36 |  |  |  |  |
| 1 | 280 | 42.86 | 4.76 | 14.29 | 38.1 | 21 | 0.0237 | 0.0732 | 0.5587 | 0.8217 |
|  | 280R | 14.71 | 23.53 | 20.59 | 41.18 | 34 |  |  |  |  |
| 2 | 280 | 52.08 | 2.08 | 18.75 | 27.08 | 48 | 0.0017 | 0.0073 | 0.5966 | 0.3135 |
|  | 280R | 21.15 | 19.23 | 23.08 | 36.54 | 52 |  |  |  |  |

In summary, these results strongly indicate that micro-pistons suspended above cells in a microchannel may be creating a static mechanical interaction and thus direct cells with respect to their shape and size by influencing elongation towards a regular alignment. Mechanical stimulation via geometric cues on cell patterning and directing cell response have also been utilized by Tse *et al*. and Hsieh *et al*., albeit in different ways to the current work [8, 9]. In the microdevices presented here, a significant percentage of the cells elongated and aligned along the radial direction, which can be assessed as directed cell behavior that can progress into a metastatic form [10]. This directedness may have been intermediated by cytoskeletal remodeling due to the change in intrinsic mechanical properties in response to the mechanical input of the micro-pistons hanging in the microchannels [26]. Although directed alignment and motility of cancer cells through solid contact guidance have been studied for potential formation of metastasis [10, 11], the mechanical impact of the hanging structures on cell cultures via no direct contact, as observed in this study, appears to require further investigation, in particular in relation to changes at protein level. In the current work, cell alignment was mainly analyzed in a label-free approach via morphological cues. As described, elongation angles of cells with respect to micropiston peripheries were measured by taking the whole cell bodies into account. This measurement method might help further the understanding of directed cell behavior through the cytoskeleton, in a manner complementary to the methods used in previous studies, which have focused more on nuclear orientation for cell angle and alignment measurements [9, 11]. By using micro-pistons of different geometries (i.e. shapes and sizes) to pattern and mechanically stimulate cells, as shown in Fig. 3 and Fig. 4, in the future, therapeutic drugs or mechanical input gradients as mechanoceuticals can be tested to attempt to distort the orientation of cells under piston peripheries.

With all observations in this study indicating that the ovarian cancer cells are able to react to the suspended structures in absence of direct solid contact, it becomes important to determine how the cells may be sensing the latter. One key difference between the main chamber (used as control) and the area under the suspended micro-pistons lies in the volume of liquid present. As such, the smaller height of the gap under micro-pistons, compared to the rest of the culture channel, may alter nutrient supply to the cells in this area and thus contribute to the observed effect. To prevent any additional stress due to nutrient factors during the static mechanical stimulation experiments, cells were kept in their cell-culture growth media at all times. Moreover, the total height of the cell-culture channel (on average 313 µm), and the gap between the bottom of the micro-piston and the glass surface (on average 108 µm) were designed to have provide appropriate geometry [27] so that cells had access to sufficient nutrients while the device was in static state. In particular, this design was expected to allow for sufficient passive diffusion of the nutrients throughout the channel.

Given the flexibility of the actuation membrane, the platform actually allows one to precisely set the gap between the micro-piston and the cells by modulating the pressure in the control channel [14]. Not only can the gap be decreased, it can also be increased, within certain constraints, by applying vacuum to the control channel. The latter leads to a retraction of the membrane, and thus the micro-piston, allowing one to take the outline beyond the on average 108 µm distance provided in the static state. The main constraint for operating the platform in this micro-piston retracted state for extended periods is the need for continuous vacuum application to the PDMS membrane. PDMS is naturally gas-permeable [28], so vacuum application may affect the composition of the culture media in the channel. Otherwise, it would be possible to use this ability to permanently retract the micro-piston to probe potential distance limits to the cell-piston interaction. Instead, in the current work we used piston-retraction only during loading to identify whether the size of the gap influenced cell behaviour. Interestingly, SKOV-3 cancer cells occupying the area under circular, triangular and square micro-pistons were observed to align across the periphery of and grow under the pistons for devices loaded via both, conventional loading, during which the piston was suspended in the middle of the culture channel in static state (Supplementary Fig. S2), and piston-retracted loading, during which the piston was retracted further from the culture channel (Supplementary Fig. S3). In future work, devices could be fabricated that contain a variety of cell-piston gaps in the static state. This would allow one to probe whether there may be a distance above which the effect is no longer observable. Biomolecular gradients could also be deliberately created within the media from the ports of the cell-culture channel towards the micro-piston within to study the effect of gradient shapes on cells [29].

To determine whether this behaviour is unique to SKOV-3 cells, micro-piston devices were also tested with other cell types including MCF7 cancer cells and MCF10A non-cancerous epithelial cells. Cell growth and alignment under the periphery of piston for these cell lines were observed to be different than for SKOV-3 cancer cells. MCF7 cells grew in clusters (Supplementary Fig. S4) and MCF10A cells moved towards each other and randomly on-chip, while forming large epithelial cell sheets (Supplementary Fig. S5). Although the three cell lines are not directly comparable, their tumorigenic and metastatic characteristics are known. Whereas MCF10As are non-tumorigenic and MCF7 tumorigenic but non-metastatic [30], SKOV-3 cancer cells are considered to be metastatic [31, 32]. Hence, their behavior in micro-piston devices was expected to be different. As detailed above, SKOV-3 cells showed response to the presence of suspended micro-piston by cell elongation and alignment patterns, and behaved in a more individual way. Conversely, MCF7 and MCF10A cells appeared to move along the chip in a more collective way, forming clusters and sheets, respectively. During this movement, MCF10A cells under micro-pistons migrated towards each other into random regions on-chip and then collectively back to under the micro-piston. This comparison suggests that different cell types can be successfully cultured in micro-piston devices and that characteristic behavior of each cell type can be captured using micro-pistons as references. Furthermore, as our observations illustrate, the device provides a promising tool even when used in static state, to differentiate the metastatic capacity of cells. However, such mechanical stimulation experiments and quantification will need to be performed with additional cell lines to confirm this capability of the device.

## 4. Conclusions

Cells are known to respond to physical cues and this plays a crucial role in tissue development and disease progression. In this paper we reported that SKOV-3 cancer cells cultured in liquid media in a cell compression microfluidic device were observed to align to geometric shapes defined by micro-piston structures suspended statically above the cell mono-layers. This effect manifested itself despite the lower faces of stationary structures being physically separated from the cells at distances of more than 30 times the height of attached cells. SKOV-3 cells aligned to square, triangular and circular micro-piston outlines and different diameters of the latter. The response of the cells to these geometries at static state was characterized for cell elongation and alignment, and compared to control areas on the same device. Results indicated statistically significant alignment for all shapes for SKOV-3 irrespective of the loading technique used, whereas no alignment was observed for control areas and MCF7 cancer cells and MCF10A non-cancerous epithelial cells. While the underlying are yet to be explained, this points to a previously undiscovered mechanotactic sensory pathway being present in SKOV-3 cells. Overall, the investigation and analysis performed here, and observation of mechanical cell response in presence of suspended micro-pistons in form of hanging structures, open up new lines of research to elaborate this and other mechanical cell stimulation effects.

## CRediT authorship contribution statement

**Sevgi Onal:** Conceptualization, Methodology, Validation, Analysis, Investigation, Writing - Original Draft, Writing - Review & Editing. **Maan M. Alkaisi:** Supervision, Writing - Review & Editing. **Volker Nock:** Conceptualization, Resources, Writing - Original Draft, Writing - Review & Editing, Visualization, Supervision, Project administration, Funding acquisition.

## Acknowledgements

We thank Helen Devereux, Gary Turner, and Nicole Lauren-Manuera for technical assistance, and Dr Tracy Hale for kindly providing us with MCF7 and MCF10A cells. The work was financially supported by the MacDiarmid Institute for Advanced Materials and Nanotechnology and the Biomolecular Interaction Centre. Additional funding for V.N. was provided by Rutherford Discovery Fellowship RDF-19-UOC-019. S.O. also thanks the University of Canterbury for a Faculty of Engineering PhD Publishing Scholarship.

**Sevgi Onal** has been a Postdoc at the Istituto Italiano di Tecnologia (IIT) in Napoli, Italy since 2023. She completed a MacDiarmid Institute sponsored PhD program in the Department of Electrical & Computer Engineering at University of Canterbury, New Zealand in 2022. Her PhD research focused on flexible microdevices for characterization of the role of bionanomechanics in cancer. She received the Master of Science degree from the Biotechnology and Bioengineering Graduate Program in the Graduate School of Science and Engineering of the Izmir Institute of Technology, Turkey in 2017. Her Master’s thesis examined the interaction mechanisms of cancer cells and macrophages on the EGF-EGFR axis by testing the hypotheses of chemotaxis, haptotaxis or direct contact in a multidisciplinary approach including microfluidics-based cell-on-a-chip technology. She has also conducted an R&D project on development of microfluidic devices with 3D printing technology in association with Initio Biomedical Ltd. Company in 2016-2017. She is a Member of the European Organ-on-Chip Society (EUROoCS). Her research interests include microfabrication, microfluidics, bionanomechanics, 3D printing, microscopy and live cell imaging.

**Maan M. Alkaisi** is an Emeritus Professor with the Department of Electrical & Computer Engineering, University of Canterbury, Christchurch, New Zealand. He is Emeritus Investigator of the MacDiarmid Institute for Advanced Materials and Nanotechnology. He is a founding member of the Nanostructure Engineering Science and Technology (NEST) research group formed back in 1998 at the University of Canterbury, which was the core group that introduced nanotechnology to New Zealand. Professor Alkaisi carried out his postgraduate studies in the UK, where he received his MSc degree from Salford University in 1976 and PhD degree from Sheffield University in 1981, both in Electronic Engineering. Professor Alkaisi is a Member of the Royal Society of New Zealand.

**Volker Nock** is a Professor and Rutherford Discovery Fellow with the Department of Electrical & Computer Engineering, University of Canterbury, New Zealand. He is a Principal Investigator and former director of the Biomolecular Interaction Center, as well as Principal Investigator of the MacDiarmid Institute for Advanced Materials and Nanotechnology and the UC Biosecurity Innovations research cluster. He received the Dipl.-Ing. degree in microsystem technology from the Institute for Microsystem Technology (IMTEK), Albert-Ludwigs University of Freiburg, Germany, in 2005, and the PhD degree in Electrical and Electronic Engineering from the University of Canterbury, New Zealand, in 2009. From 2009 to 2012, he was a MacDiarmid and Marsden Research Fellow. His research interests focus on microfabrication, materials, and microfluidic devices at the interface of biology, chemistry, and medicine. He is a member of the Royal Society of New Zealand, microSANZ and senior member of IEEE.

