## Supplemental Information for "On-chip non-contact mechanical cell stimulation - quantification of SKOV-3 alignment to suspended microstructures"

#### Supplementary Methods

**Table S1.** Media formulations used for the culture of SKOV-3, MCF7 and MCF10A, including base media type and concentrations of the supplements.

| Medium for | Component | Final Concentration | Stock Concentration | Volume to add |
|---|---|---|---|---|
| SKOV-3 | MEM (L-Glu positive) | - | - | 500 ml |
| | Fetal Bovine Serum | 10% | 100% | 56.3 ml |
| | Pen-Strep | 1X | 100X (10,000 U/ml) | 5.63 ml |
| | Fungizone | 0.2% | 100% | 1.126 ml |
| Cell dissociation | 0.05% Trypsin-EDTA, no phenol red | 1X | 100X | - |
| | Add 1X PBS or Gibco DPBS, no calcium, no magnesium to dilute Trypsin-EDTA and wash cells | | | |
| MCF7 | DMEM high glucose pyruvate (L-Glu positive) | - | - | 450 ml |
| | Fetal Bovine Serum | 10% | 100% | 50 ml |
| | Insulin Human Recombinant Zinc Solution | 10 µg/ml | 4 mg/ml | 1250 µl |
| | Pen-Strep | 1X | 100X (10,000 U/ml) | 5 ml |
| Cell dissociation | 0.25% Trypsin-EDTA, phenol red | - | - | - |
| | Add DMEM high glucose complete media to stop action of trypsin, then Gibco DPBS, no calcium, no magnesium for washing cells | | | |

| | | | | |
|---|---|---|---|---|
| MCF10A | DMEM/F12 (L-Glu positive) | - | - | 500 ml |
| | Horse Serum | 5% | 100% | 25 ml |
| | Human EGF | 20 ng/ml | 100 µg/ml | 100 µl |
| | Hydrocortisone | 0.5 µg/ml | 1 mg/ml | 250 µl |
| | Cholera Toxin | 100 ng/ml | 1 mg/ml | 50 µl |
| | Insulin Bovine Pancreas | 10 µg/ml | 10 mg/ml | 500 µl |
| | Pen-Strep | 1X | 100X (10,000 U/ml) | 5 ml |
| Cell dissociation | 0.05% Trypsin-EDTA, phenol red | - | - | - |
| | Add resuspension media DMEM/F12 (L-Glu positive) supplemented with 20% (final) Horse Serum and 1% Pen-Strep to stop action of trypsin and for washing cells | | | |

### Supplementary Figures

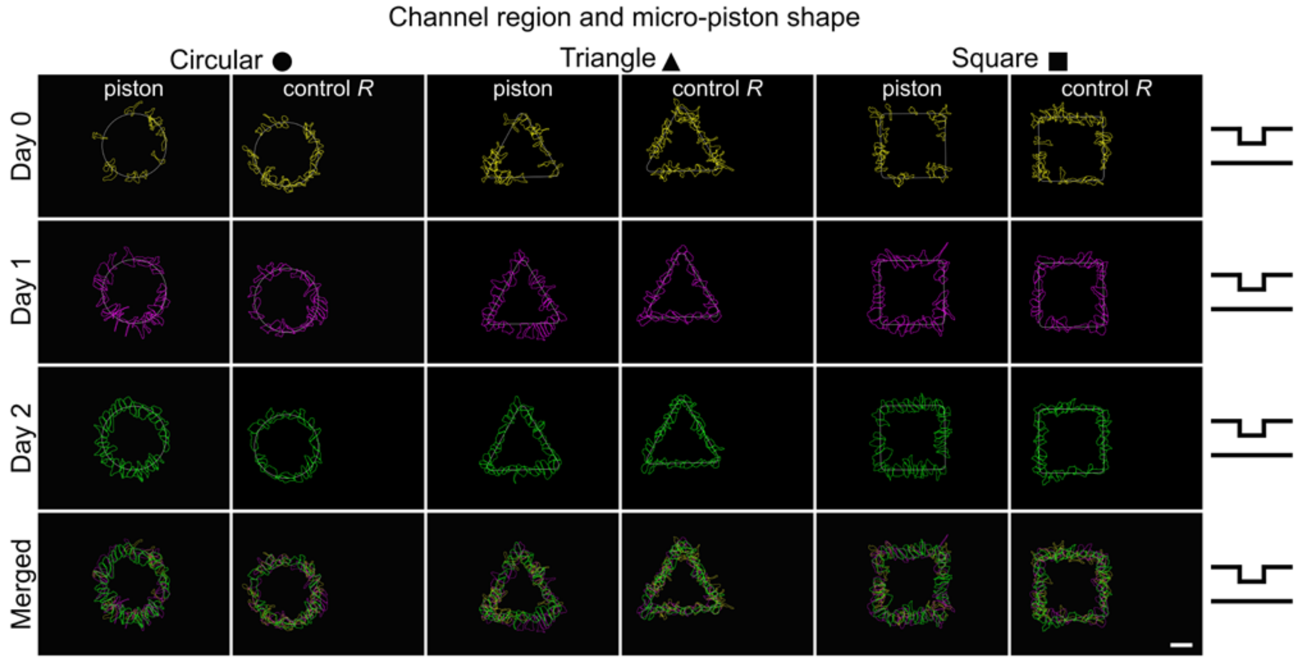


**Figure S1:** Cell alignment profiles under the micro-piston periphery and in the control region, showing separate and overlaid outlines of cells over days with respect to the micro-piston shape being circular, triangular or square, color-coded as yellow for Day 0, magenta for Day 1, green for Day 2 (Scale bar 100 µm).

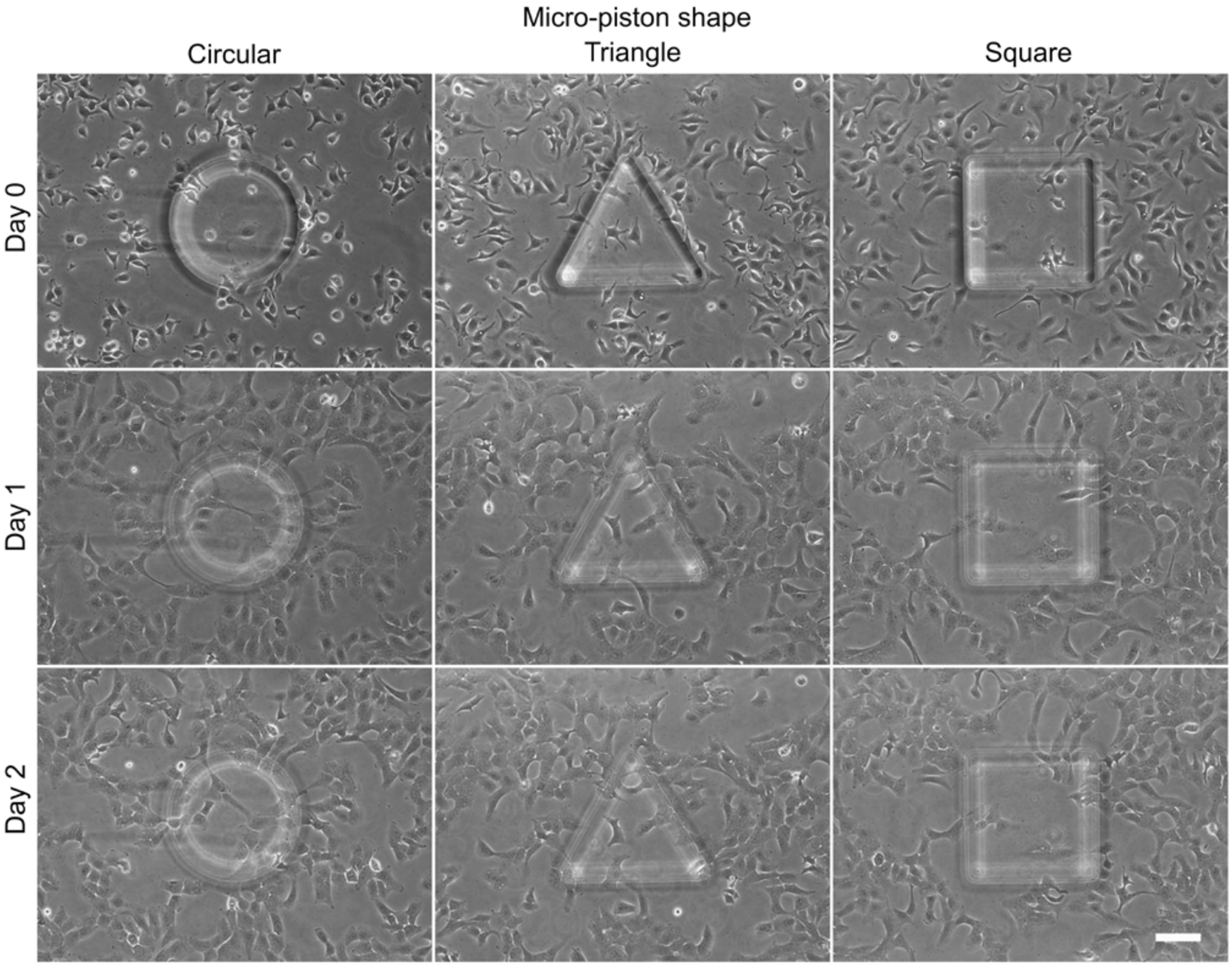


**Figure S2:** Representative images for SKOV-3 cancer cells occupying the area under circular, triangular and square micro-pistons in devices loaded with cells by conventional loading during which the piston was suspended in the middle of the culture channel in static state. Cells were observed to align across the periphery of and grow under the pistons (Scale bar 100 µm).

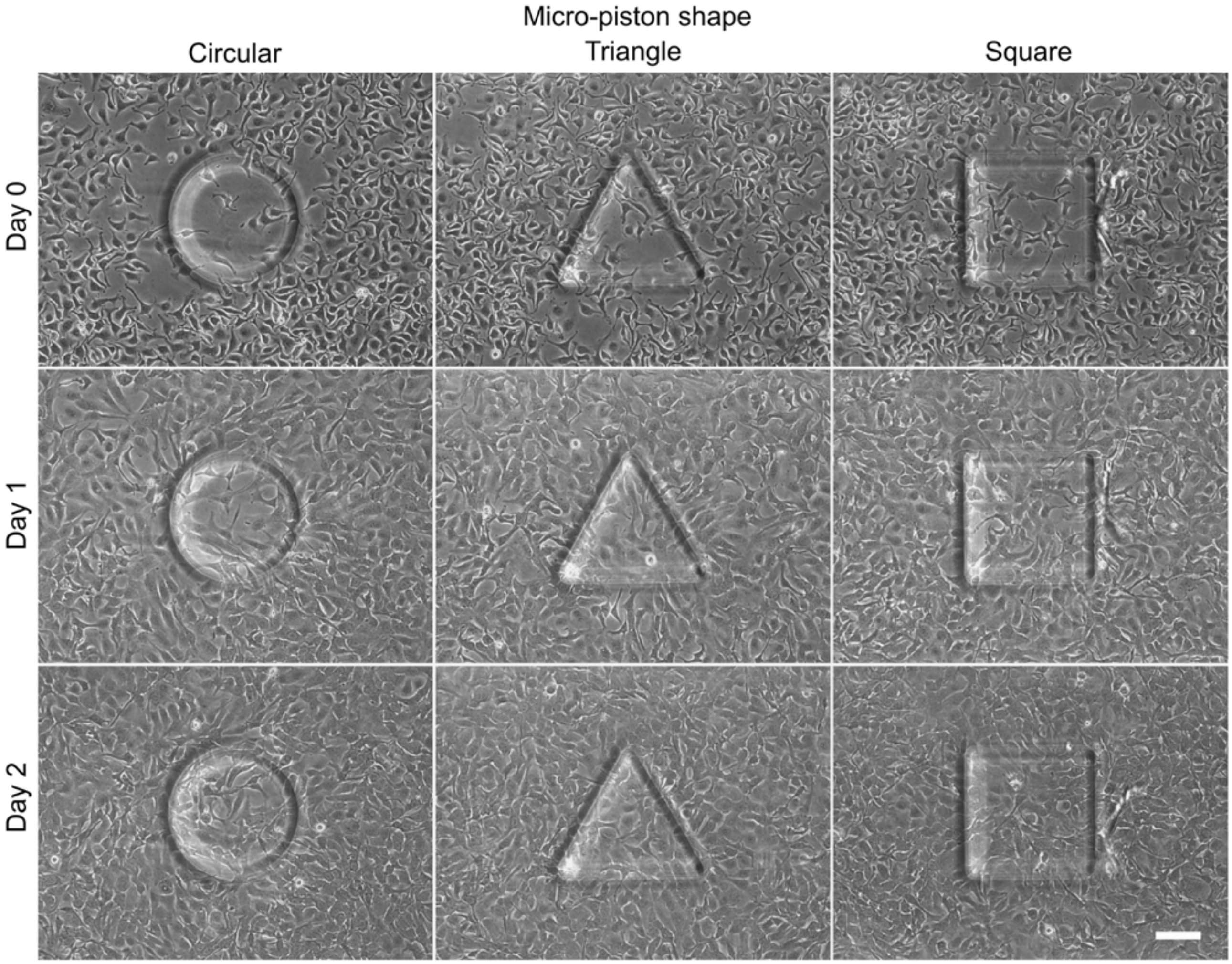


**Figure S3:** Representative images for SKOV-3 cancer cells occupying the area under circular, triangular and square micro-pistons in devices loaded with cells by piston-retracted loading. Cells were also observed to align across the periphery of and grow under the pistons (Scale bar 100 µm).

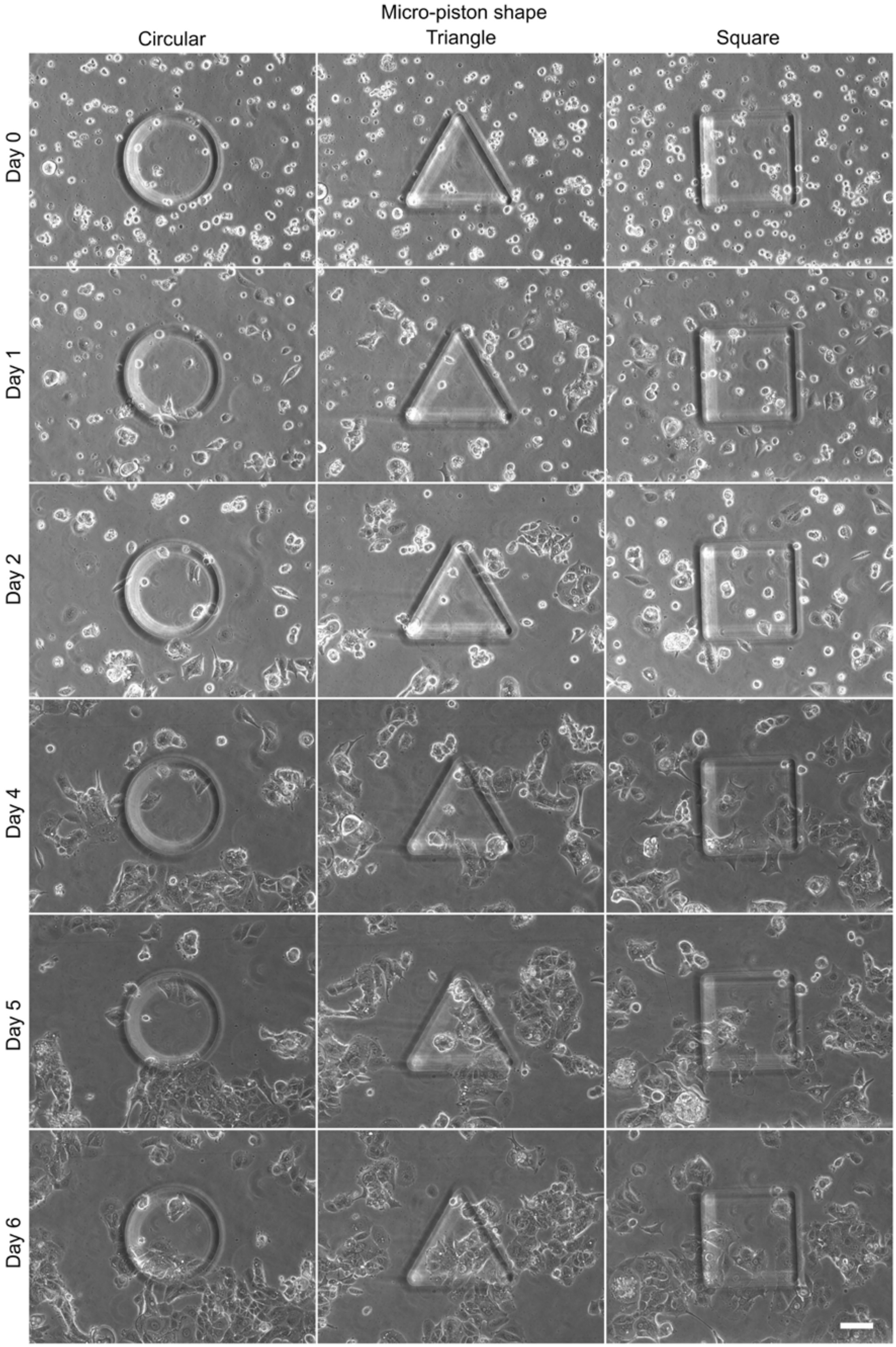


**Figure S4:** Representative images for MCF7 cancer cell behavior on-chip around and under circular, triangular and square micro-pistons in devices loaded with cells by piston-retracted loading (Scale bar 100 µm).

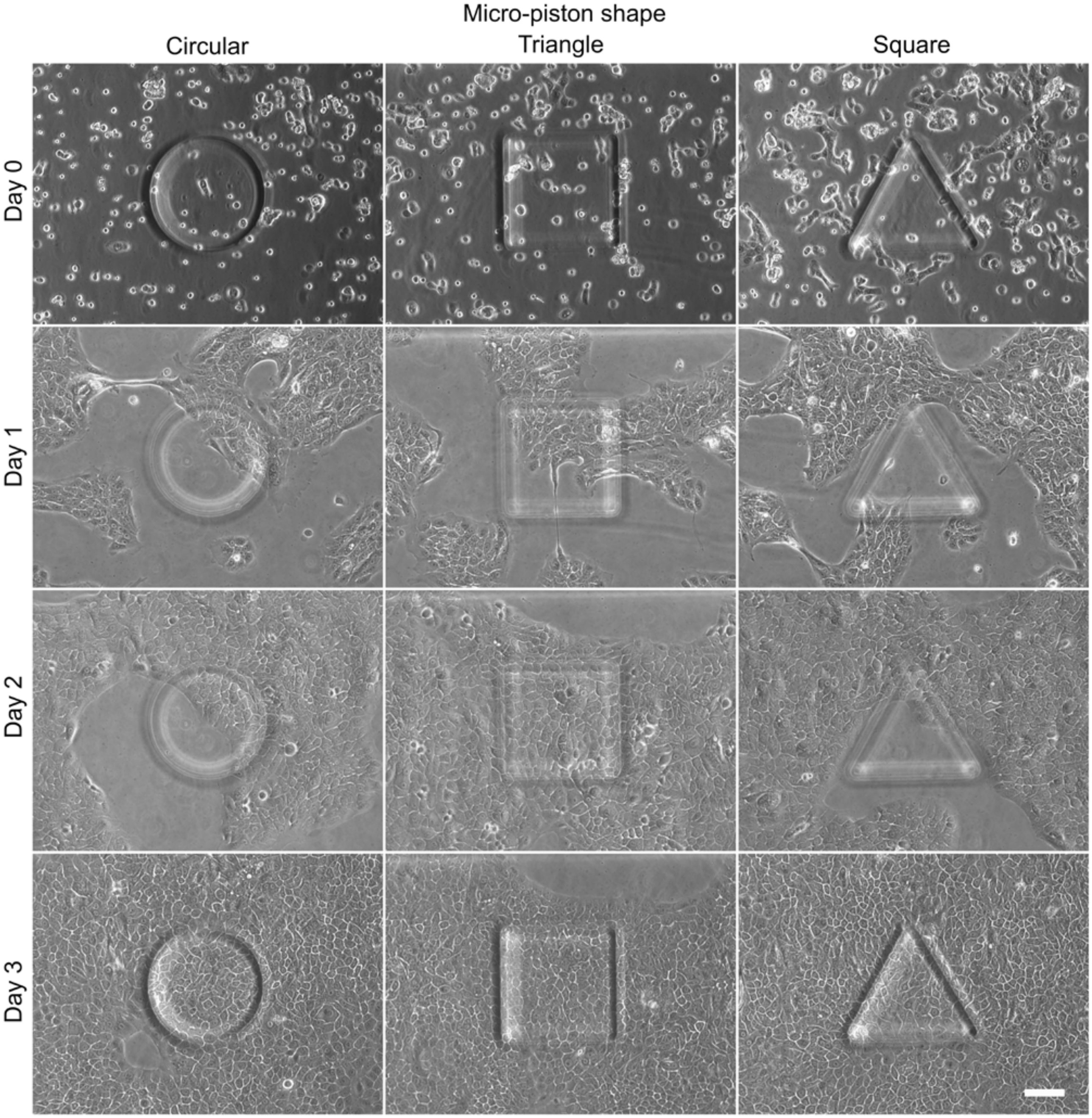


**Figure S5:** Representative images for MCF10A epithelial cell behavior on-chip around and under circular, triangular and square micro-pistons in devices loaded with cells by piston-retracted loading (Scale bar 100 µm).

#### Supplementary Movies

**Movie S1.** Bright-field microscopy live-cell image sequences (CytoSmart) of SKOV-3 cancer cells. Before and after 48-hours of live imaging, phase-contrast images were acquired using an inverted microscope. Cell outlines were drawn using freehand tool of ImageJ (Fiji), which show cell elongation and alignment with respect to the periphery of the micro-piston. **Clip 1:** SKOV-3 cells proliferating inside a PDMS device with 300 µm square micro-piston and orienting under the periphery of the suspended micro-piston over a period of 48 hours. Images were recorded at every 30 minutes in an incubator of 37°C and 5% $CO_2$. Automated analysis of area coverage is shown as inset. **Clip 2:** SKOV-3 cells proliferating inside a PDMS device with 300 µm triangular micro-piston and orienting under the periphery of the suspended micro-piston over a period of 48 hours. Images were recorded at every 30 minutes in an incubator of 37°C and 5% $CO_2$. Automated analysis of area coverage is shown as inset. **Clip 3:** SKOV-3 cells proliferating inside a PDMS microchannel of around 313 µm height without a micro-piston inside over a period of 48 hours. Images were recorded at every 30 minutes in an incubator of 37°C and 5% $CO_2$. **Clip 4:** SKOV-3 cells proliferating inside a PDMS microchannel of 200 µm height without a micro-piston inside over a period of 48 hours. Images were recorded at every 30 minutes in an incubator of 37°C and 5% $CO_2$.
